# Rapid metagenomic sequencing for diagnosis and antimicrobial sensitivity prediction of canine bacterial infections

**DOI:** 10.1101/2023.01.30.526267

**Authors:** Natalie Ring, Alison S. Low, Bryan Wee, Gavin K. Paterson, Tim Nuttall, Richard Mellanby, David Gally, J. Ross Fitzgerald

**Author notes:** **Corresponding author and email address** J. Ross Fitzgerald. **Repositories:** All raw sequencing data (nanopore) produced during this work have been deposited under NCBI BioProject Accession PRJNA925092 (https://www.ncbi.nlm.nih.gov/bioproject/PRJNA925092).

## Abstract

Antimicrobial resistance is one of the greatest current threats to human and animal health. There is an urgent need to ensure that antimicrobials are used appropriately to limit the emergence and impact of resistance. In the human and veterinary healthcare setting, traditional culture and antimicrobial sensitivity testing is typically conducted, requiring 48-72 h to identify appropriate antibiotics for treatment. In the meantime, broad-spectrum antimicrobials are often used, which may be ineffective or impact non-target commensal bacteria. Here, we present a rapid diagnostics pipeline, involving metagenomic Nanopore sequencing directly from clinical urine and skin samples of dogs. We have optimised this pipeline to be versatile and easily implementable in a clinical setting, with the potential for future adaptation to different sample types and animals. Using our approach, we can identify the bacterial pathogen present in a sample with 100% sensitivity within 5 hours. For urine samples, we can predict antibiotic sensitivity with up to 95% accuracy. However, skin swabs which exhibited lower bacterial abundance and higher host DNA, were less amenable and an additional host depletion step may be required prior to DNA extraction. In summary, our pipeline represents an important step towards the design of individually tailored veterinary treatment plans on the same day as presentation, facilitating effective use of antibiotics and promoting antimicrobial stewardship.

**Impact statement:** Antimicrobial resistance (AMR) is a major threat to veterinary and human healthcare. It is a one-health problem, as humans and dogs are in close contact, require similar antibiotics, and share bacterial pathogens and AMR genes. Limited treatments options due to AMR would have a catastrophic effect. The risk of infection would render much of modern healthcare (including critical care, orthopaedic and complex surgeries, implants and oncology) impossible. In addition, routine infections could become life threatening. It is therefore critical to preserve the efficacy of these drugs for the future. Inappropriate antimicrobial use is the single biggest factor driving AMR. Antimicrobial stewardship involves reducing antimicrobial use, using first-line narrow-spectrum drugs, and avoiding overly long treatment. Delays in culture-based diagnosis lead clinicians to speculatively use broad-spectrum antibiotics and prolong courses of treatment beyond clinical cure. Our rapid diagnostic approach will have a major impact in reducing, refining and replacing antibiotic use. This will advance antimicrobial stewardship in veterinary and human healthcare.

**Data summary:** All sequencing data mentioned in this work is available from NCBI, BioProject PRJNA925092, Biosamples SAMN32880396 to SAMN32880438, run accessions SRR23195371 to SRR23195413.

**The authors confirm all supporting data, code and protocols have been provided within the article or through supplementary data files**.

## Introduction

Antimicrobial resistance (AMR) levels amongst human and veterinary bacterial pathogens are escalating globally, to the extent that the World Health Organization now classifies AMR as one of the biggest threats to global health, food security and development (1, 2). The current gold-standard diagnostic methods used in veterinary practice are culture-based or involve remote service providers for PCR, requiring several days to yield results. Broad-spectrum or inappropriate antibiotics are often started while waiting for the results; this inappropriate antimicrobial use is a major driver for AMR (3). Furthermore, animals with suspected highly contagious or potentially zoonotic infections like leptospirosis may require quarantining while waiting for a diagnosis leading to escalated costs, wasted resources, and unnecessary stress for the quarantined animal (4).

Novel methods that enable sensitive, same-day rapid diagnosis and prediction of antibiotic sensitivity across different infection types are required. One method which is of particular interest is metagenomic whole genome sequencing (WGS), involving the extraction of all genomic DNA (gDNA) present in a sample and subsequent identification of pathogens by unbiased DNA sequencing. One of the main sequencing platforms for this kind of rapid diagnosis is the MinION, produced by Oxford Nanopore Technologies (ONT), which has been tested with clinical sputum, endotracheal aspirate, blood, urine and other sample types (5-13). Importantly, the MinION’s small size and relative affordability could enable its usage in a wide variety of clinical settings. In addition, sequence data is accessible in real-time as it is produced, reducing the time required to identify pathogens and their AMR genotypes to as little as 10 minutes (14, 15). The long reads produced by nanopore sequencing readily facilitate whole genome assembly which potentially allows the linkage of AMR genes to specific bacterial strains (16).

Here, we set out to develop a rapid, culture-free metagenomic sequencing pipeline to identify pathogens and predict AMR in canine samples in a veterinary hospital setting. To initially develop this pipeline, we chose to focus on two common canine infections: urinary tract infections (UTIs) and skin infections (pyoderma). Antibiotic therapy is often the first line of treatment in these infections, but AMR, including multi-drug resistance, is frequently observed, and increasing in prevalence (17-21). Furthermore, recurrent infections are common, leading to frequent return visits to the clinic and further courses of antibiotics (22, 23). A rapid, sensitive method for diagnosing the bacterial pathogens and predicting their antimicrobial sensitivity could therefore prevent the use of inappropriate antibiotics, and limit the amount of clinical care required. Although we focussed on urine and skin swab samples here, we aimed to design a protocol which could be adapted to an array of other sample types (e.g. blood) from infections in other animals including humans. Moreover, the approach could be used in a variety of clinical settings, from small practices to large hospitals. By comparing and optimising a number of different kit-based gDNA extractions and sequencing techniques, combined with community-built DNA analysis tools, we have developed a pipeline which can identify the bacterial species present in clinical samples with around 100% sensitivity and specificity, in as little as 5 hours. It can also predict the antimicrobial resistance phenotype of those species with up to 95% accuracy, in around 8 hours.

## Methods

### Selection of DNA extraction kit

Three different kits recommended in the literature were tested: DNeasy Blood + Tissue (Qiagen, Hilden, Germany), DNeasy Powersoil (Qiagen) and MagAttract HMW DNA (Qiagen). To test each kit, overnight *Escherichia coli* CAN-50 growth in Luria-Bertani (LB, ThermoFisher, Massachusetts, USA) broth equivalent to 10^9^ CFUs was spun down at 16,000 *xg*, and the cell pellet was resuspended in 1 ml healthy dog urine. The urine-cell suspension was then processed using each kit, according to the relevant manufacturer’s instructions for each, finishing with an elution into 50 µl nuclease-free water. The resulting extractions were used to compare the kits in terms of (i) gDNA yield in 50 µl nuclease-free water (quantified by Qubit dsDNA HS kit), (ii) lysis method for Gram-positive species (enzymatic vs bead-beating), (iii) speed, and (iv) cost.

### Optimisation of metagenomic bacterial lysis

Records for canine urine and skin swab samples processed at the Royal (Dick) School of Veterinary Studies Hospital for Small Animals (HfSA) in Edinburgh between 2018 and 2019 were analysed to determine which species were most commonly detected, so the broad efficacy of our extraction protocol for the most relevant pathogens could be tested (**Supplementary Table S1**). Metapolyzyme (Sigma-Aldrich, Missouri, USA), an enzymatic lysis cocktail containing six lysis enzymes (achromopeptidase, chitinase, lyticase, lysostaphin, lysozyme and mutanolysin), was trialled for our extraction protocol. An isolate of the Gram-positive species *Staphylococcus pseudintermedius*, ED99 (24), was used to trial four enzymatic lysis options: lysostaphin, lysozyme, and two different concentrations of metapolyzyme. Cells were grown overnight on tryptic soy agar (TSA, Oxoid ThermoFisher, Massachusetts, USA) plates at 37°C, then a single colony was transferred into tryptic soy broth (TSB, Oxoid) media and cultured overnight at 37°C with shaking. 2.5 ml of overnight culture was pelleted by centrifuging for 3 min at 16,000 *xg*, then resuspended in 3 ml healthy dog urine.

For each of the four lysis options tested, the resulting urine-and-cell suspension was centrifuged for 3 min at 16,000 *xg*, then resuspended in 160 µl 50 mM Tris, 10 mM EDTA, pH 8.0 (buffer P1 for the MagAttract HMW DNA kit). 20 µl lysozyme (100 mg ml^-1^), lysostaphin (10 mg ml^-1^), metapolyzyme (6.6 mg ml^-1^) or metapolyzyme (3.3 mg ml^-1^) was added, and the solution mixed by flicking. The solution was then incubated on a thermomixer for 1h at 37°C with 900 RPM shaking. After 1h, 20 µl proteinase K was added, and the solution was incubated for a further 30 min at 56°C with 900 RPM shaking. The rest of the MagAttract HMW DNA Gram-positive protocol was then followed as per the manufacturer’s instructions (starting from step 8 on page 26 in the MagAttract HMW DNA Handbook 03/2020), eluting into 50 µl nuclease-free water as the final step. DNA concentrations were quantified using the Qubit dsDNA HS kit according to the manufacturer’s instructions.

### Testing our extraction protocol on different species

The optimised MagAttract + Metapolyzyme extraction protocol was tested on the most commonly isolated species identified in the HfSA’s records: *E. coli, S. pseudintermedius, Streptococcus canis, Enterococcus faecalis, Pseudomonas aeruginosa, Proteus mirabilis, Pasteurella canis, Klebsiella pneumoniae, Kocuria kristinae/Kocuria rosea*, and *Clostridium perfringens*. For aerobic species, cells were grown on TSA plates at 37°C for 24h or 72h (*Kocuria/Streptococcus*), then in TSB medium at 37°C with shaking for 24h or 72h (*Kocuria*/*Streptococcus*). For two anaerobic or facultative anaerobic species (*C. perfringens* and *P. canis*), cells were grown on TSA plates at 37°C in a sealed box with Anaerogen sachets (Oxoid ThermoFisher) for 24h, then in TSB medium in growth flasks in a sealed box with Anaerogen sachets at 37°C with shaking for 24h. For all species, 3 ml of broth culture was pelleted by centrifuging at 16,000 *xg* for 3 min, the pellet was resuspended in 3 ml healthy dog urine, then pelleted again in the same way. The pellet was then processed as per the optimised protocol, starting with resuspension in 160 µl buffer P1 and 60 minutes lysis with 20 µl metapolyzyme. DNA was eluted into 50 µl nuclease-free water, and quantified using Qubit’s dsDNA HS kit.

### Quality of the extracted DNA

The purity of the extracted DNA was assessed using a Nanodrop spectrophotometer (ThermoFisher) to measure the 260/280nm and 260/230nm absorbance ratios of 22 clinical samples (indicated in **Supplementary Table S2**). The samples were then cleaned up using the ProNex Size-Selective Purification System (Promega, Wisconsin, USA) according to manufacturer’s instructions, starting with 50 µl DNA and 200 µl ProNex beads, and eluting into 20 µl nuclease-free water at the end of the protocol. The purity of the cleaned-up DNA was then measured again by Nanodrop.

### Final optimised protocols for metagenomic DNA extraction and clean-up

The final optimised protocols, including post-extraction clean-up with ProNex beads, are available from protocols.io:

dx.doi.org/10.17504/protocols.io.n2bvj8o5bgk5/v2 (urine)

dx.doi.org/10.17504/protocols.io.q26g7yr19gwz/v2 (skin swabs)

### Species identification strategies

For each experiment detailed here, species were identified from sequencing reads using either ONT’s EPI2ME tool (v3.0.1-7052513, https://epi2me.nanoporetech.com/, ONT account required) or Kraken2 (v2.1.1, 25) with one of two custom databases, “pathogens_plus” or “bacteria_plus”. The bacteria_plus database was constructed from all bacterial representative genomes present in the NCBI RefSeq database in November 2022 (4,032 species) plus eight mammalian genomes (*Canis lupus familiaris, Homo sapiens, Felis catus, Equus caballus, Oryctolagus cuniculus, Sus domesticus, Bos taurus* and *Ursus arctos*). The “pathogens_plus” database contains 668 genomes of various bacterial, viral, protozoan and fungal pathogens, including the top 100 European human and animal pathogens identified in a 2014 study (26), plus selected other pathogens known to be important in veterinary samples, such as *Leptospira* and *Mycobacterium spp* (a full list of species included can be seen in **Supplementary Table S3**). The pathogens_plus database also contains the same eight mammalian genomes included in the bacteria_plus database. The tool and database used is noted for each step of the protocol development below.

### Optimising the sequencing protocol

Two rapid library preparation kits were tested: rapid PCR barcoding (SQK-RPB004, Oxford Nanopore Technologies (ONT), Oxford, UK) and rapid barcoding (SQK-RBK004, ONT). The SQK-RPB004 PCR reaction was carried out on 22 clinical samples (**Supplementary Table S2**), according to the manufacturer’s instructions, and the concentration of each was measured by Qubit’s dsDNA HS kit before and after the reaction.

All 22 samples prepared with the rapid PCR barcoding kit were subsequently sequenced on MinION R9.4.1 flow cells (ONT) according to the SQK-RPB004 protocol, although ProNex beads were substituted into steps which required AMPure XP beads. Four of these samples, two urines (DTU09 and DTU16) and two skin swabs (SkSw08A and SkSw14), were re-sequenced with the rapid barcoding kit, to compare the results of sequencing with and without the PCR amplification. The DNA was prepared according to the manufacturer’s instructions for SQK-RBK004, except for the substitution of ProNex in place of AMPure XP again. The results of the two library preparation methods were compared in terms of (i) run yield, and (ii) bacterial reads vs eukaryotic host contamination (according to ONT’s online EPI2ME classification “WIMP” tool).

#### Determining the lower detection limits for the rapid barcoding (SQK-RBK004) kit

Three sequencing runs were conducted using serially diluted gDNA extracted from *E. coli, S. pseudintermedius* and *S. canis*, respectively. For each, a starting sample of gDNA was serially 1:1 diluted in nuclease-free water, from around 10 ng µl^-1^ down to below 0.1 ng µl^-1^. Each dilution was prepared for sequencing with SQK-RBK004 as described above, with a different barcode used for each dilution and barcode01 used as a negative control (nuclease-free water). The dilution series (plus negative control) from each species was then sequenced on a fresh MinION R9.4.1 flow cell for 24 h. The sequencing results for each dilution were compared in terms of (i) run yield, and (ii) reads from the expected species vs background contamination (according to EPI2ME).

#### MinION vs Flongle flow cells

Two relatively highly concentrated gDNA samples were used to compare MinION flow cells with Flongle flow cells: a clinical skin swab sample (SkSw21, 55 ng µl^-1^) and a 1:1 mix of *E. coli* and *P. aeruginosa* isolate DNA, both extracted previously during the DNA extraction optimisation trials (80 ng µl^-1^). Each sample was prepared for sequencing on an R9.4.1 Flongle flow cell according to the Flongle-specific manufacturer’s instructions for SQK-RBK004 (using ProNex beads instead of AMPure XP), then each was sequenced separately for a full 24h on the Flongle device, and the total yield of the 24 h run was noted. More DNA from the same samples was then prepared for sequencing on an R9.4.1 MinION flow cell with SQK-RBK004 as above. Sequencing was started for the first (*E. coli + P. aeruginosa*) sample on a Mk1C MinION device, and the sequencing was monitored. The sequencing was stopped when the run yield matched that of the 24 h Flongle run, and the time taken to reach that yield was recorded. The MinION flow cell was then washed using the flow cell wash kit (EXP-WSH004, ONT) according to the manufacturer’s instructions, and the second sample (SkSw21) was loaded and sequenced. Again, the sequencing was monitored closely and stopped once the run yield matched that of the 24 h Flongle run.

#### Determining the optimal MinION flow cell usage strategy

Two flow cells and a variety of DNA samples were used to test how many times a MinION flow cell can be washed and re-used. For the first flow cell, the DNA sample was a relatively high concentration previously extracted *E. coli* isolate (110 ng µl^-1^) which had both been stored at 4°C since extraction. An initial sample of 7.5 µl of DNA was prepared for sequencing with SQK-RBK004 as previously described, using barcode01. Sequencing commenced on a fresh flow cell and was stopped when a target of 200 Mbp was reached. The time taken to reach 200 Mbp was recorded, and the flow cell was washed using EXP-WSH004 as previously described. A second 7.5 µl was processed in the same way, with barcode02, followed by another wash with EXP-WSH004. This protocol continued until the time taken to reach 200 Mbp was longer than 2 h, using the next subsequent barcode (barcode03, barcode04, etc.). For each sequencing step, the starting number of available sequencing pores was recorded at the beginning of the run. After each run, the sequenced reads were quality controlled by EPI2ME and NanoStat (v1.6.0, 27), and the percentage of DNA assigned to the wrong barcode was noted.

For the second flow cell, the same protocol was followed, but freshly extracted DNA from clinical samples, ranging from 0.76 to >120 ng µl^-1^, was sequenced. Each sample was sequenced for up to 3 h, and some samples were sequenced simultaneously (with different barcodes). Full details of the samples sequenced are given in **Supplementary Table S4**.

#### Testing adaptive sampling on a GridION to reduce contamination with host DNA

Two flow cells and two DNA samples were used to test the efficiency of adaptive sampling for reducing levels of host DNA in clinical samples. The two DNA samples, one an *E. coli* isolate and the other a sample of previously sequenced canine DNA, were 75 ng µl^-1^ each. The samples were mixed in various ratios (90:10, 75:25, 50:50, 25:75 and 10:90). 15 µl of each mix, plus 15 µl of a previously sequenced clinical sample known to be contaminated with host DNA, were prepared for sequencing with SQK-RBK004 (volumes doubled), each with a different barcode, and using nuclease-free water with barcode01 as a negative control. Half of the pooled library was then sequenced on a fresh R9.4.1 flow cell on a GridION, with real-time super accuracy basecalling and no adaptive sampling. The other half was sequenced at the same time on a second fresh R9.4.1 flow cell on the GridION, with real-time super accuracy basecalling and adaptive sampling selected to deplete DNA which mapped to a canine reference genome provided to the software (GCA_014441545.1 ROS_Cfam_1.0).

The resulting datasets for each barcode were analysed using Kraken2 (v2.1.1, 25) with the bacteria_plus database described above. The percentage of reads from each sample assigned to *Escherichia* (or *Shigella*) and *Canis lupus familiaris* were recorded, and the differences between the percentages of each with and without adaptive sampling were tested for significance using a paired Wilcoxon signed-rank test in R (v4.1.2).

### Testing clinical samples

#### Sample selection, DNA extraction and sequencing, and flow cell usage

During the development of this protocol, DNA from 45 clinical urine and skin swab samples was extracted and sequenced, using either the rapid barcoding or rapid PCR barcoding library preparation kit (**Supplementary Table S3**). These samples were not usually processed on the same day they were received; instead, they were processed and sequenced in batches, and the raw sample and/or extracted DNA was stored at 4°C in the meantime. After these development steps, a further nine urine samples were processed in batches of one to four. These were collected from the HfSA and processed immediately using the finalised extraction protocol (https://www.protocols.io/view/magattract-metapolyzyme-metagenomic-gdna-extractio-chnrt5d6).

The extracted DNA was prepared for sequencing using the rapid barcoding kit (SQK-RBK004) as described above, using barcode 01 for a negative control (nuclease-free water, which was processed using the same extraction protocol as the real samples) and the remaining sequential barcodes for the real samples. Samples were sequenced for 2h, or until 100 Mbp had been sequenced, whichever was sooner. In between sequencing runs, the flow cell was stopped, washed using the flow cell wash kit (EXP-WSH004) according to manufacturer’s instructions and stored with storage buffer at 4°C until the next use.

#### Species identification and AMR prediction

During sequencing, the data produced underwent preliminary analysis using EPI2ME’s Fastq Antimicrobial Resistance workflow. After sequencing, the data were further analysed, including species identification with Kraken2 (pathogens_plus database), genome assembly with Flye (v2.9-b1774, 28) using the --meta flag and no --genome-size flag, genome annotation using Prokka (v1.14.5, 29) using the appropriate --species and --genus flags along with --compliant, and --usegenus, and AMR prediction using Abricate (v1.0.1, 30) with the NCBI database downloaded on 18 January 2022 and AMRFinderPlus (v3.10.45, 31) with the database version 2022-10-11.2.

During development, large volumes of data were sequenced for some samples. For the analysis described above, these large datasets were randomly down-sampled to 100 Mbp using Rasusa (v0.6.1, 32). The results from all data analyses were collated to predict which species had been present in the original sample, and the AMR phenotypes expected from them. In addition, the length of time taken for each sample was recorded. The results from this pipeline were then compared to the results from the Veterinary Pathology Unit (VPU) at the HfSA, which were produced using the current gold-standard techniques of culture followed by species identification and antibiotic susceptibility testing (AST) using a VITEK® 2 (bioMérieux) instrument.

## Results and Discussion

### Comparison of DNA extraction methods

A wide variety of DNA extraction methods are available, and many of them have been used in previous studies aiming to develop rapid diagnostic protocols or to extract metagenomic DNA from urine samples (7, 9, 11, 14, 16, 33-37). We selected three commonly used kits to test here (**Table 1**), based on their previous successful application and their potential flexibility for application to other sample types in the future. A metagenomic approach using enzymatic lysis rather than mechanical cell disruption should result in high molecular weight (HMW) fragments, facilitating better genome assembly and more efficient species and AMR gene identification. Nonetheless, we tested one kit using mechanical disruption, to compare the efficiency of mechanical versus enzymatic lysis.

**Table 1:** gDNA extraction kits tested, and their pros and cons

| Kit | Pros | Cons | Yield from test *E. coli* extraction (µg) | Cost per sample (kit only) |
| --- | --- | --- | --- | --- |
| **Qiagen DNeasy Blood + Tissue** | • Quick extraction (<40 minutes after lysis)<br>• Easy, column-based<br>• No additional reagents or equipment required<br>• No bead-beating | • Different Gram +ve protocol (enzymatic lysis step required) | 3.16 | £3.92 |
| **Qiagen DNeasy PowerSoil** | • Very quick extraction (30 minutes or less)<br>• Same protocol for Gram +ve or -ve, optimised for metagenomics extractions | • Extremely low yield<br>• Bead-beating, highly sheared DNA<br>• Not optimised for liquid samples | 0.6 | £6.06 |
| **Qiagen MagAttract HMW DNA** | • Optimised for extraction of HMW DNA<br>• No bead-beating<br>• Magnetic beads and 6 wash steps, so theoretically very clean DNA even from dirty starting samples<br>• Quick extraction (30 minutes after lysis) | • Different Gram +ve protocol (enzymatic lysis step required)<br>• Magnetic rack required | 3.08 | £4.75 |

Of the three kits we trialled, the one with mechanical lysis (the DNeasy Powersoil) was the most expensive per-sample, and produced lowest yield of gDNA when tested (Table 1). The two kits that included optional enzymatic Gram-positive lysis steps were the DNeasy blood + tissue kit and the MagAttract HMW kit, each producing similar yields of DNA in a similar time-frame when tested with *E. coli*. Although the spin column-based DNeasy kit had a lower cost-per-sample, we ultimately selected the MagAttract in our protocol due to its optimisation towards extracting HMW DNA, and its extensive washing steps that should limit the presence of inhibitors that affect library preparation.

### Our optimised protocol can efficiently extract DNA from the bacterial pathogens most commonly identified in canine urinary tract and skin infections

We examined records from the HfSA to identify the array of bacterial species most commonly associated with clinical urine or skin swab samples from dogs (**Figure 1, Supplementary Figure S1**). Overall, 90% of culture-positive cases were associated with ten different genera, with a total of 53 different genera in total, including 48% Gram-positive and 52% Gram-negative (**Supplementary Table S1**). Thus, we required a lysis method that would allow the extraction of DNA from a wide variety of different species. Metapolyzyme contains six enzymes optimised for the lysis of bacterial and/or fungal cell walls. Of these six, lysostaphin, mutanolysin and lysozyme should lyse all of the Gram-positive species most commonly seen in the 2018-2019 HfSA data. We determined that 3.3 mg ml^-1^ of metapolyzyme resulted in better yield of extracted DNA than either lysozyme or lysostaphin for the most common Gram-positive species in skin swabs (*Staphylococcus pseudintermedius*, **Table 2**). The optimised MagAttract + Metapolyzyme extraction protocol was next tested on the top ten species identified in urine and/or skin swab samples (**Figure 1, Table 3**). Each species was grown in appropriate media and spiked into 3 ml healthy dog urine to simulate a clinical sample, and our protocol was successful in extracting micrograms of DNA from all ten species, including anaerobes, Gram-positives, and slow-growing bacteria (e.g. *Kocuria*). We therefore concluded that the protocol would be efficacious in extracting DNA from the vast majority, if not all, of the species encountered in clinical canine urine or skin swab samples.

**Table 2:** Different lysis enzymes tested with *S. pseudintermedius* ED99 DNA extraction

| Lysis enzyme | DNA yield (µg) |
| --- | --- |
| Lysozyme (100 mg/ml) | 0.46 |
| Lysostaphin (10 mg/ml) | 2.75 |
| Metapolyzyme (6.6 mg/ml) | 5.5 |
| Metapolyzyme (3.3 mg/ml) | 5.2 |

**Table 3:**
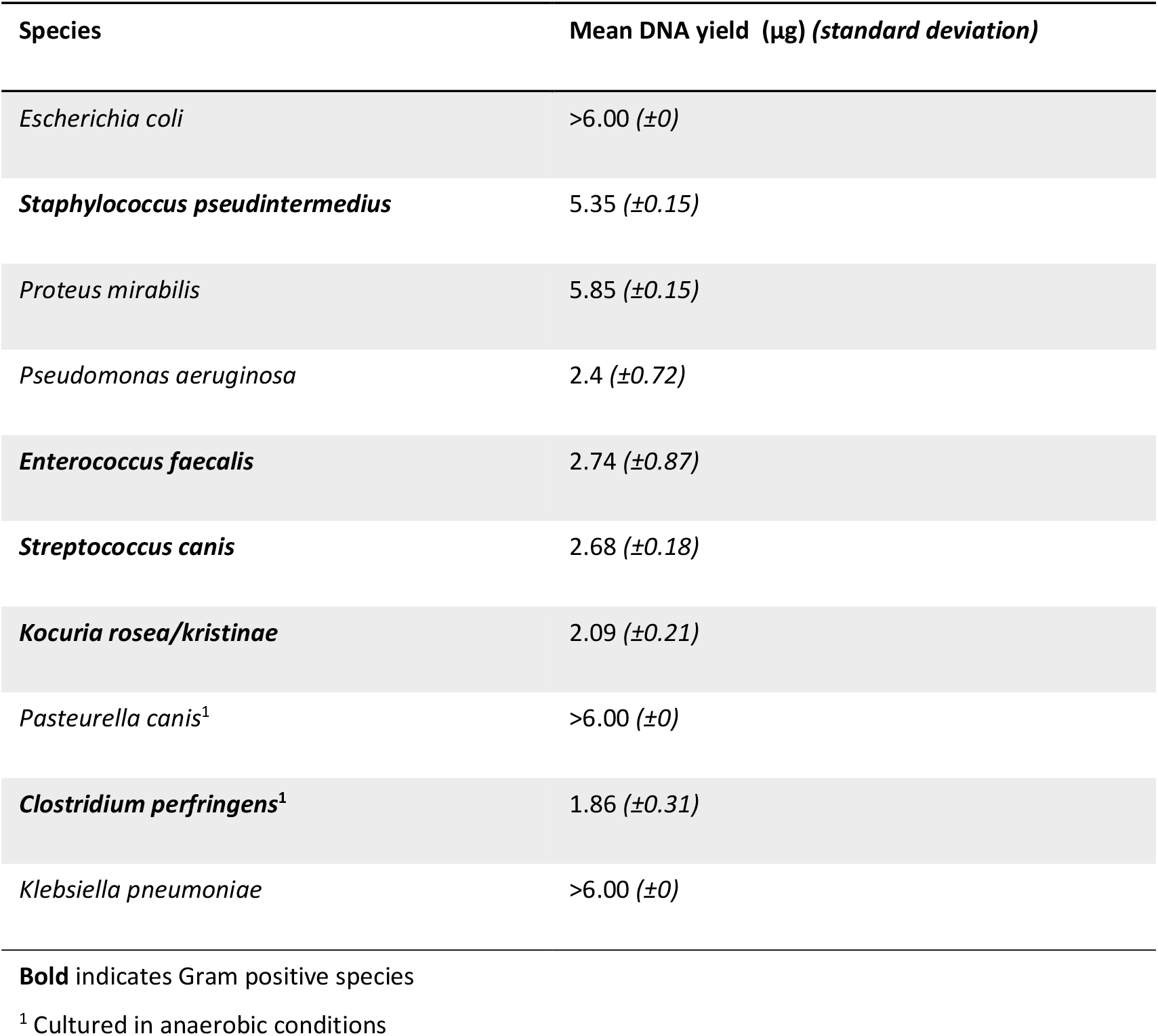
gDNA concentrations extracted from ten most commonly encountered species, using our optimised lysis and extraction protocol

**Fig. 1.**
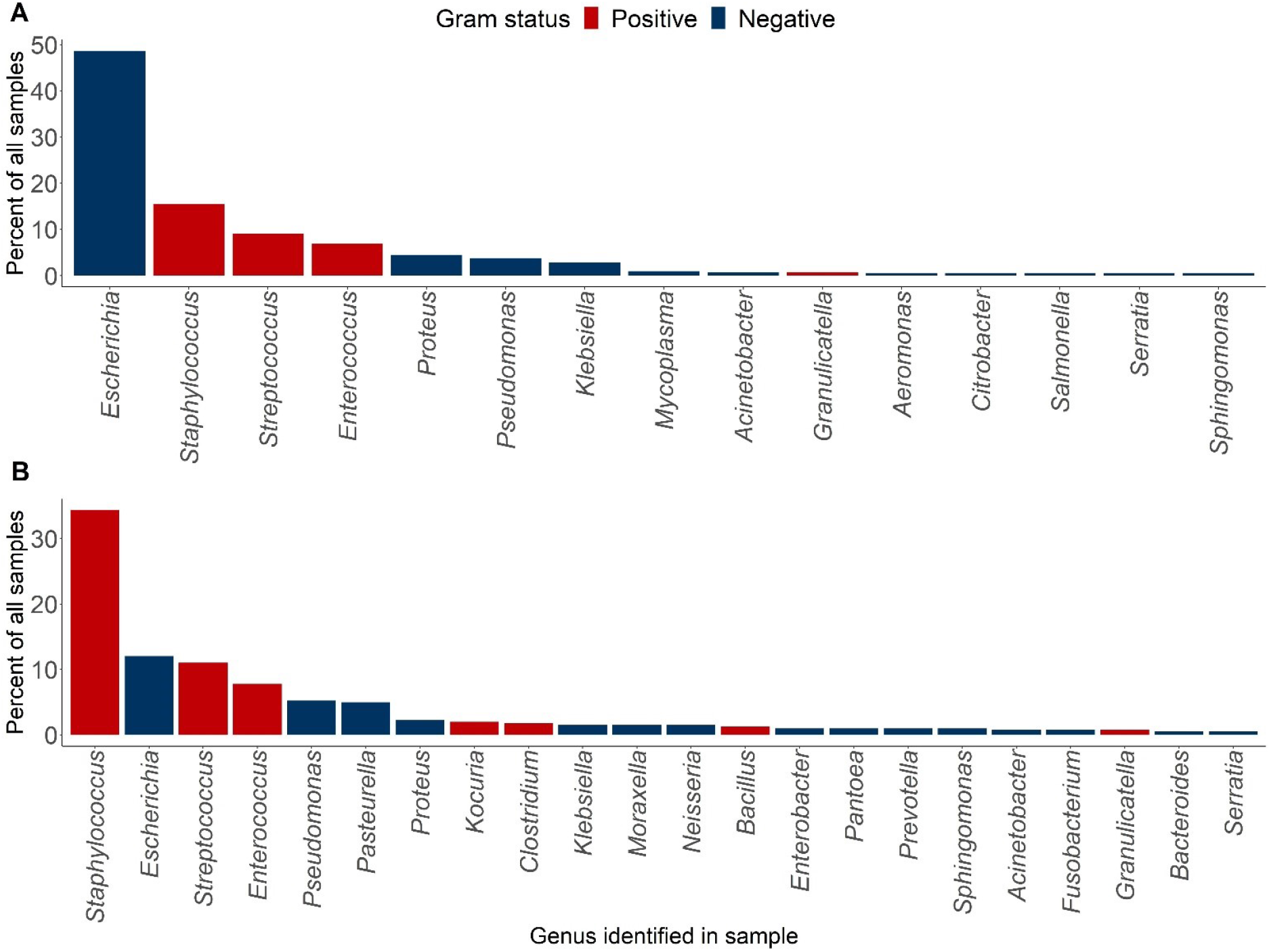
Pathogens identified in HfSA (A) urine and (B) skin swab samples, 2018 & 2019. Species observed in only one sample (17 species for urine samples, 22 for skin swab samples) are not shown here, but can be seen in **Supplementary Table S1**.

### Enhancement of DNA purity during extraction

As clinical samples may contain contaminants or inhibitors that impact library preparation or sequencing, we tested if a further clean-up step would be enhance the protocol. Nanodrop 260/280nm and 260/230nm absorbance ratios were used to assess the purity of 22 clinical samples (14 urine, 8 skin swabs, **Supplementary Figure S2**) before and after a clean-up protocol using the ProNex Size-Selective Purification System. This kit was selected because it should maintain the high molecular weight of the DNA extracted by the MagAttract protocol, and for its affordability relative to the AMPure XP system. After the clean-up, the mean value across the 22 samples was close to ideal at 1.82, although the difference between pre- and post-clean-up ratios was not assessed to be significant according to the Wilcoxon signed rank test for paired samples (*p*=0.7, *n*=22). In contrast, the 260/230nm ratios were significantly improved post-clean-up (*p*=0.004, *n*=22); the mean value of 1.36 was not optimal but still enabled efficient library preparation. Our findings support the inclusion of the additional clean-up step between DNA extraction and library preparation, despite adding extra time to the final length of the rapid diagnostics protocol. An additional benefit of the clean-up protocol is that by eluting into a final volume lower than the starting volume (20 µl vs 50 µl) the DNA is further concentrated.

### Rapid barcoding enables sequencing and identification of species from extracted concentrations of DNA as low as 0.04 ng µl^-1^

Our experience extracting DNA from the first few clinical samples (**Supplementary Table S2**) indicated that DNA concentrations may be low in some samples (the mean concentration of the first five skin swab samples was just 4.86 ng µl^-1^, while the mean from the first five urine samples was 21.98 ng µl^-1^). Some previous protocols developed for rapid bacterial identification by whole genome nanopore sequencing have used the rapid PCR barcoding library preparation kit, SQK-RPB004 (7, 9, 38, 39). The major benefit of this kit is the amplification of DNA during the PCR step, which may be important for low abundance samples, though with the disadvantage of the additional time required. However, we found that this step resulted in unpredictable yields of DNA and a tendency to amplify host DNA (**Supplementary figures S3 and S4)**. We therefore trialled the rapid barcoding kit without the PCR step (SQK-RBK004) instead, aiming to determine the lowest concentration which could be sequenced and still produce enough usable data to identify selected bacterial species from our samples. We conducted three serial dilutions of DNA samples extracted from cultured *E. coli, S. pseudintermedius* and *S. canis*, and found that the relevant bacterial species was identifiable at concentrations much lower than our means of 4.86 (skin swabs) and 21.98 (urine) ng µl^-1^. For *E. coli* (**Figure 2B**) and *S. canis* (**Figure 2D**), the original species could be detected above background contamination even at the lowest concentrations tested (0.07 and 0.04 ng µl^-1^. For *S. pseudintermedius* (**Figure 2C**), the original species could be detected above background contamination at 0.16 ng µl^-1^. We therefore concluded that we could use the rapid barcoding kit to sequence our clinical samples without the need for the PCR amplification step.

**Fig. 2.**
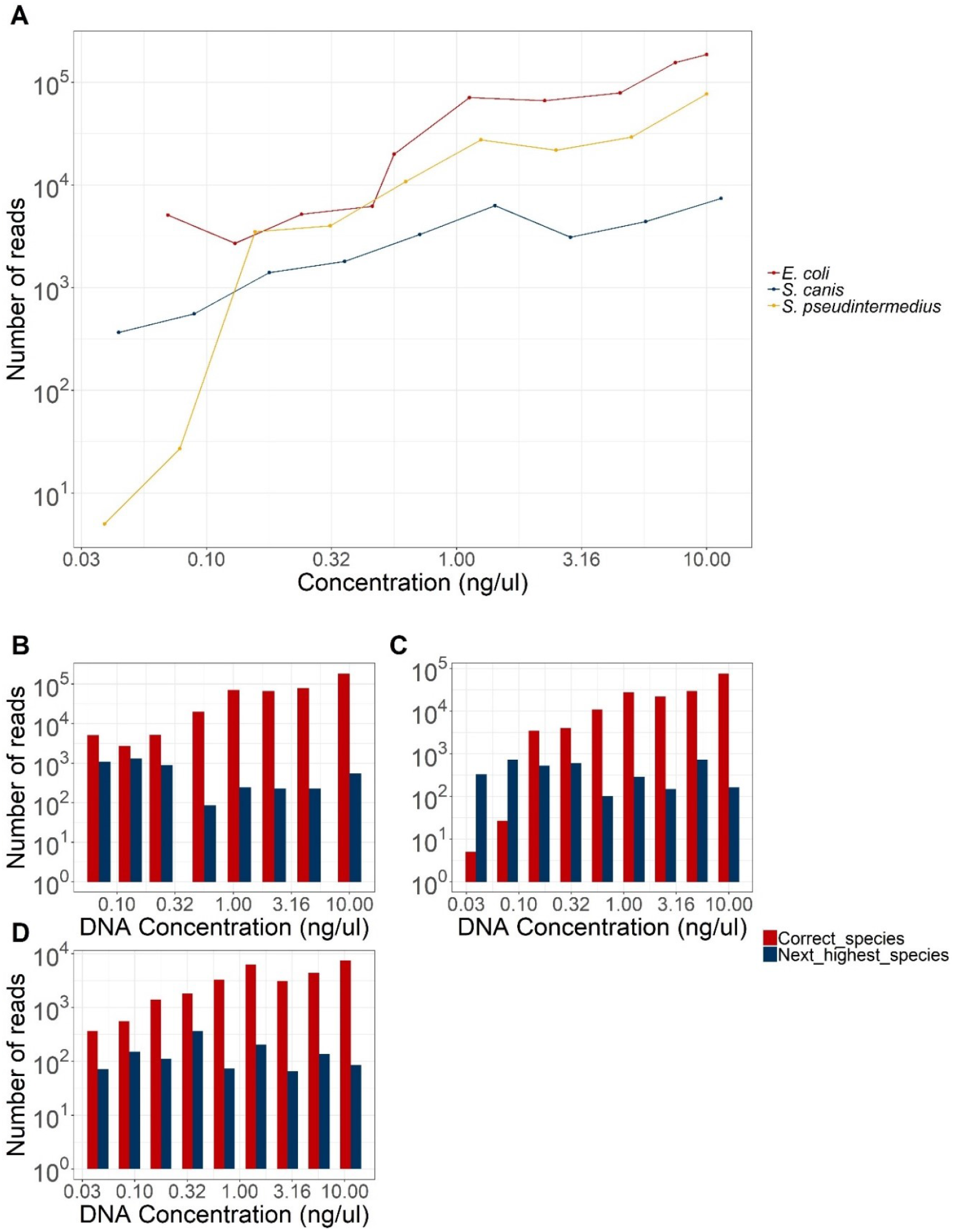
Sequencing serial dilutions of *E. coli, S. pseudintermedius* and *S. canis*. 1:1 dilution of gDNA in nuclease-free water, sequenced for 24 hours using SQK-RBK004 (rapid barcoding) and R9.4.1 MinION flow cells. A) shows the number of reads identified by EPI2ME’s WIMP tool as the correct species vs starting gDNA concentration. B), C) and D) show the number of reads identified as the correct species vs the number identified as the next most common species for *E. coli, S. pseudintermedius* and *S. canis*, respectively.

### Flongle flow cell use and re-use of MinION flow cells

The Flongle is an adapter which fits into MinION or GridION sequencers and allows the use of Flongle flow cells, which are a single-use, lower yield alternative to MinION flow cells. These characteristics of Flongle flow cells are desirable for clinical use: single-use means no potential cross-contamination between different samples, and the lower yield means the flow cells are correspondingly less expensive than MinION flow cells. Clinical applications for which Flongle flow cells are already being used include rapid sequencing of viruses such as SARS-CoV-2 and monkeypox (40, 41), HLA-typing (42, 43), and 16S metagenomics (44, 45). A small number of previous studies have also investigated the use of Flongle flow cells for the rapid identification and typing of bacterial infections (5, 46, 47).

We sought to evaluate the utility of Flongle flow cells in our protocol compared to the classical MinION using two clinical samples (**Supplementary Table S4**). The volume of DNA sequence data produced by Flongle in a 24h period was matched by MinION within 1.5h for one sample, and 3h for the other. However, although MinION offers greater sequencing speed, the price of each MinION flow cell precludes the use of one flow cell per sample. Barcoding allows multiplexing of up to 12 samples, but in a clinical setting, there may not be sufficient samples to load a full flow cell and still produce results within the desired timeframe. We therefore examined the possibility of reusing MinION flow cells. This would involve sequencing a sample for as long as necessary to produce the sequence data required, stopping the run and performing a DNase wash of the flow cell, then storing the flow cell until another sample was received. In this way, the same flow cell could theoretically be used many times, thus reducing the cost-per-sample without the need for simultaneous sequencing of multiple samples. Accordingly, we aimed to establish i) how many times could we re-use a flow cell to produce sufficient sequence data in a timely manner, and ii) how much residual DNA from previous samples would remain in the flow cell after washing. Employing a cultured *E. coli* sample, we examined the capacity to produce 200 Mbp of DNA within 2h, and what percentage of the reads had the correct barcode attached. We were able to use the same flow cell eight times before it was exhausted, and the percentage of reads with the correct barcode never fell below 98.8% (**Figure 3, Supplementary Table S5**). Further, using different barcodes for each subsequent sample reduces the risk of cross-contamination between runs to negligible. We also later decided that, for most samples, only 100 Mbp of sequence will be sufficient for species identification and AMR prediction, hence flow cells could potentially be used even more times.

**Fig. 3.**
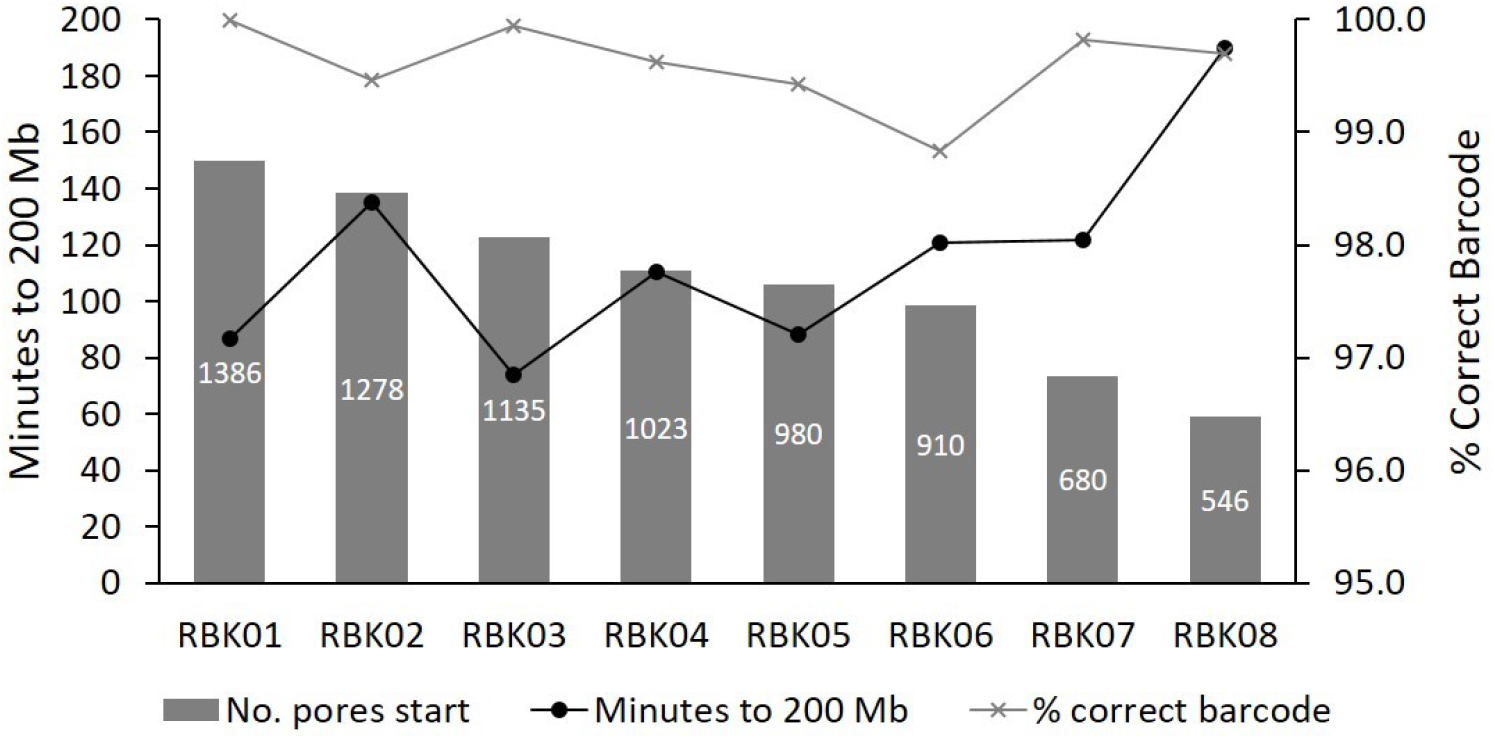
MinION flow cell progression when washing and re-using multiple times. The same 108 ng µl^-1^ *E. coli* sample was sequenced up to 200Mbp in 8 consecutive sequencing runs on a single flow cell. In between runs, the sequencing was ceased, and the flow cell was washed with wash kit EXP-WSH004-XL according to manufacturer’s instructions. The flow cell was either then stored over one or two nights with storage buffer, or the next sequencing run was immediately commenced.

Importantly, we also trialled a second flow cell, using real clinical urine samples that had been extracted on the same day they were sequenced (**Table 4**). This second flow cell was able to produce data for nine samples before being used to exhaustion, with samples being sequenced in batches of one to four per run, and over a period of roughly one month.

**Table 4:**
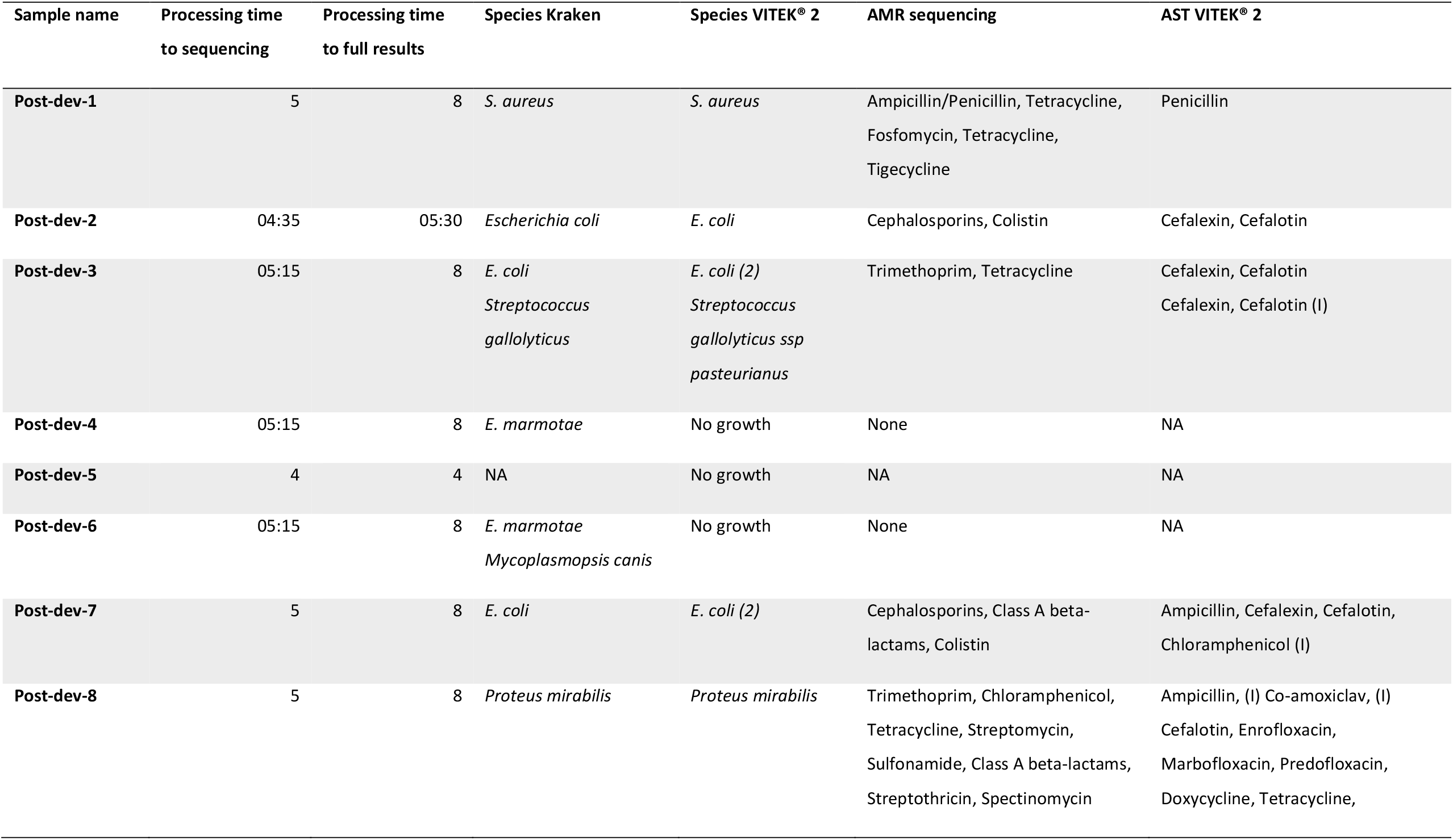
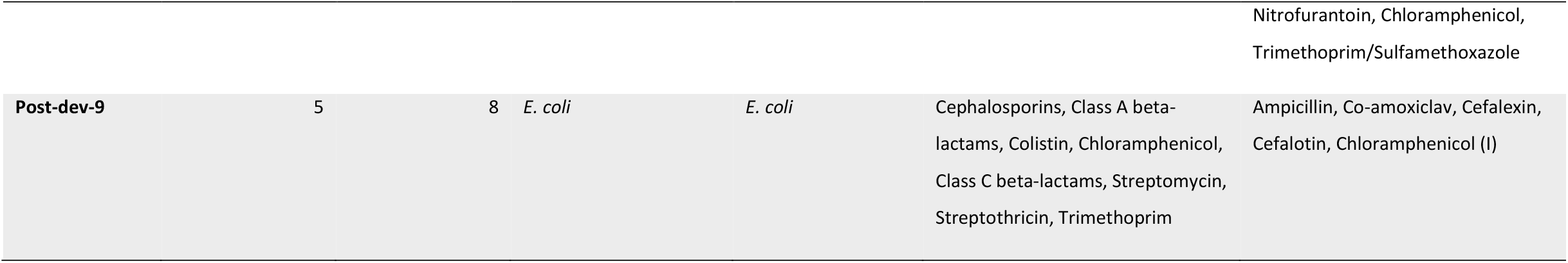
Results from processing 9 clinical urine samples in real-time using our final optimised protocol

Ultimately, we estimate that the average MinION flow cell can be used and re-used around eight times to sequence 100-200 Mbp per sample. This slight increase in cost is balanced against producing the data at least eight times faster than it would be on a single-use Flongle flow cell, as well as producing less plastic waste (especially as MinION flow cells can be returned to ONT for recycling, whilst Flongle flow cells cannot).

### Adaptive sampling on a GridION significantly reduced host DNA contamination, but not enough to benefit sample analysis

The pipeline developed here is primarily aimed at processing clinical samples and it is therefore likely that many of the DNA samples extracted will contain host DNA alongside the bacterial DNA. In some cases, such as purulent skin swabs or urine samples containing high numbers of white blood cells, the vast majority of the DNA extracted may be derived from the host, which could impact on the accuracy of the diagnostic and AMR prediction from bacterial DNA. Lab-based methods to reduce the numbers of host cells in the sample prior to DNA extraction exist, but can be costly and/or slow, so were not included in this pipeline. As an alternative, a sequencing-based sampling method was trialled.

Adaptive sampling is technique unique to nanopore-based sequencers, in which DNA reads are compared to a reference genome whilst the strand of DNA is still being sequenced. The technique can be used either to enhance or to deplete sequences which map to the reference genome. Due to the computer power needed, MinION sequencers are unlikely to be capable of adaptive sampling, unless connected to a GPU-powered computer. The GridION sequencer, however, is equipped with GPU processors, and adaptive sampling is therefore possible.

Although adaptive sampling did significantly reduce the proportion of eukaryotic reads (*p*=0.03135, *n*=6) and increase the proportion of bacterial reads (*p*=0.03135, *n*=6) in our paired samples, the absolute differences in percentages were small: 1% fewer eukaryotic reads, and 0.7% more bacterial reads (**Supplementary Table S6**). These differences would realistically have little effect on our ability to identify bacterial species or detect AMR genes. Where a GridION is available, we would therefore recommend its use in our protocol, but where only a MinION is available, the ability to produce accurate results will not be affected.

### Our optimised protocol produces up to 100% sensitivity and specificity for species prediction, within five hours

During the development of this protocol, DNA was extracted from 45 urine and skin swab samples (20 urines and 25 skin swabs). Subsequently, we processed a further nine urine samples using the final optimised protocol, for a total of 54 clinical samples tested (**Supplementary Tables S2, S7, and S8, and Table 4**).

A wide variety of DNA concentrations were retrieved from these samples, ranging from 0 to >120 ng µl^-1^ for the urine samples and 0 to 116 ng µl^-1^ for the skin swabs (**Figure 4**). As mentioned previously, the lower limit of detection for sequencing was determined to be < 0.1 ng µl^-1^ of cleaned-up DNA. In total, only five urine samples and five skin swabs produced DNA with lower concentrations than our lower limit and colony forming units (CFU) per ml of sample indicated that those samples had either no growth of any kind, or contained fewer than 1×10^5^ cells. We therefore estimate our lower limit for sufficient DNA from clinical samples to be between 1×10^5^ and 1×10^6^ CFU/ml. The exact lower limit is likely to depend on other sample characteristics, such as the presence of host cells, sediment or inhibitors.

**Fig. 2.**
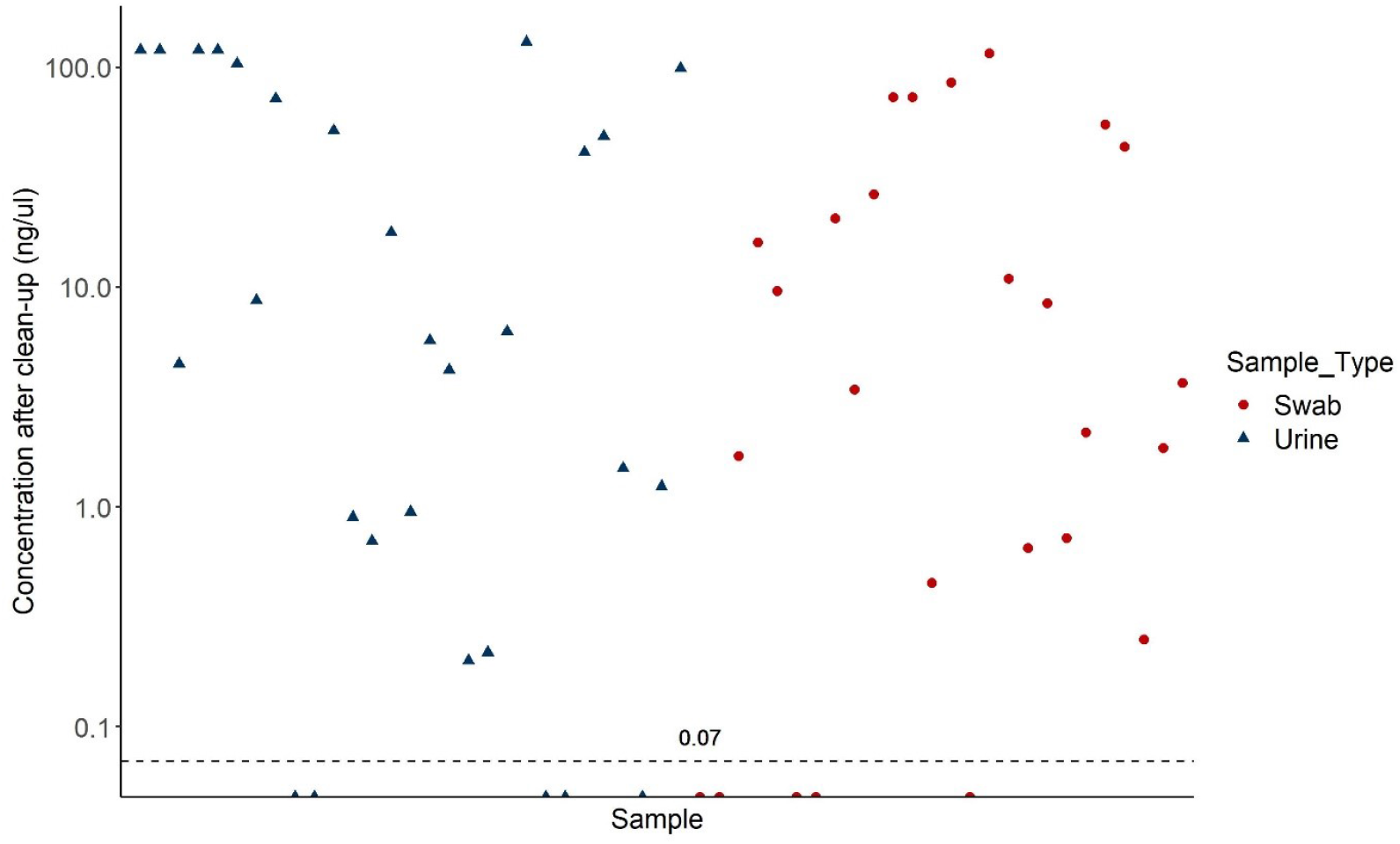
Concentration of DNA extracted from 54 real skin swab and urine samples from the HfSA. DNA was extracted using our optimised extraction protocol, followed by clean-up with ProNex beads and quantification with the Qubit HS DNA kit. Nine samples fell below the lower limit of detection for sequencing (0.07 ng μl-1)

In order to test relevance to a clinical scenario, the nine post-optimisation urine samples were processed in real-time. The average time taken to process the samples through to the commencement of sequencing was just under 5 hours (**Table 4 and Supplementary Table S8**). Using the online EPI2ME tool, species identification was carried out in real-time as soon as the first sequencing reads were produced. The speed of EPI2ME can be affected by the number of concurrent online users, but we were able to produce preliminary species identification calls within approximately 20 min of sequencing. In each case, we continued to sequence until 100 Mbp of reads had been produced and, to reduce the likelihood of false positive calls, we also performed species identification using Kraken2 after sequencing was complete. Of note, the initial rapid EPI2ME species calls differed from the Kraken2 calls for only one sample in which EPI2ME detected both *P. mirabilis* and *E. coli*, whilst Kraken2 only identified *P. mirabilis* at an abundance greater than 1% of the reads. The culture-based diagnostic process also only identified *P. mirabilis*. Across all the clinical samples processed during the study, EPI2ME and Kraken2 agreed on the species present in all samples except this one, and one other (where EPI2ME detected *Finegoldia magna* in addition to *E. coli* and *E. faecalis*, but Kraken2 did not). Likewise, the culture-based identification for the samples were in almost 100% agreement with both EPI2ME and Kraken2 (**Table 4 and Supplementary Table S8**). For 2 samples, SkSw8A and SkSw10, our sequence-based protocol revealed additional species according to EPI2ME (Kraken2 only identified an additional species for SkSw10). Although these may be false-positives, it is also highly feasible that the sequence-based approach is be more sensitive than the current gold-standard culture-based techniques. It is worth noting that while culture-based species identification and ASTs are “gold standard” they are themselves not 100% accurate, with problems including small sample sizes compared to the infecting population, results skewed to easily cultured and fast growing organisms, and laboratory error (e.g. single doubling dilution steps between sensitive, intermediate and resistant). A 2007 evaluation estimated VITEK® 2 accuracy for species identification to be 98.3%, while AST accuracy was estimated to be 97.7% (48).

### Metagenome-based AMR prediction is dependent on bacterial sequence data volume

The accuracy of using sequencing data alone for AMR identification varies from species to species, but previous studies have suggested >97% accuracy for *E. coli* and over 99% for *Staphylococcus aureus* (49, 50). AMR phenotypes can be predicted from sequencing data alone, by screening the data for known AMR-related genes or SNPs, based on curated databases like ResFinder or CARD (51, 52). This level of accuracy is, however, dependent on the volume of sequencing data available, the complexity of the samples being sequenced, and the amount of contaminating host DNA present. Here we interrogated DNA sequence produced from both urine samples and skin swabs using our metagenomic protocol for AMR resistance determinants and compared to the phenotypic data produced by AST (**Supplementary Tables S2, S7 and S8, and Table 4**).

For urine samples, 71.7% of the resistance phenotypes identified by phenotypic AST were also predicted by our protocol (*n*=53). 50% of samples were in exact agreement for all AMR calls for each sample (*n*=16). However, further investigation revealed that the vast majority of AST calls that did not correlate with the sequence-based predictions were defined as intermediate resistance, often to chloramphenicol, which may represent unknown genetic mechanisms. Excluding intermediate resistance calls from the VITEK® 2 results, we detected 83.7% of the same AMR using sequencing data alone. Excluding a single missed gene in one of our post-optimisation samples (post-dev-8) increases the sensitivity of our AMR predictions to 95.3% (41 of 43 resistant phenotypes accurately predicted, excluding intermediates).

The remaining two AMR phenotypes not detected by our pipeline were from one sample (post-dev-3), which was determined by the VITEK® 2 to be a co-infection consisting of two different strains of *E. coli* and one strain of *Streptococcus gallolyticus ssp pasteurianus*. Though the type of resistance (multiple cephalosporins) identified in this sample was frequently detected accurately by our pipeline, the complexity of this sample may have hindered our AMR prediction, and we suggest that complicated co-infection samples may require significantly more than 100 Mbp of sequencing data, particularly when host DNA is also present (Kraken2 identified 47.5% of the DNA in this sample as canine). Taken together, our pipeline can predict the vast majority of resistant phenotypes in non-complex urine samples.

In contrast, for skin swab samples, of which 5 had also been phenotyped by AST, effective sequence-based prediction of AMR was not possible due to the low abundance of the bacteria in the samples, (3×10^2^ to 7.5×10^5^ cfu/ml), almost all below our predicted lower limit of detection. In addition, four of the five samples were more than 95% dog DNA, so bacterial DNA comprised a very low proportion of the extracted DNA (reflecting 1 or 2x coverage). Thus, for low abundance skin samples, species can easily and quickly be identified by sequencing, but the amount of sequencing data required to accurately detect AMR genotypes is likely cost-prohibitive for veterinary applications, as it would reduce the number of uses of each flow cell. One of our urine samples (post-dev-9) contained over 83% dog DNA, as well as a multidrug-resistant *E. coli*, yet we were able to predict all four of its non-intermediate AMR phenotypes from just 100 Mbp. This suggests that even small increases in the relative levels of bacterial cells we are seeing in our low abundance skin swabs could greatly improve our ability to accurately detect AMR genotypes. Although we have shown that removing host DNA via adaptive sampling has limited benefit, host-depletion steps could be added prior to DNA extraction to enable more accurate AMR prediction even in lower abundance infections. Previous studies have shown promising results for host depletion using a range of techniques including saponin-and-DNase enzymatic methods, PMA plus UV-light-based chemical methods, and even methods as simple and cost-effective as physical filtering of host cells through a 22 µm filter prior to extraction (9, 53, 54).

### Concluding comments and future considerations

We have developed and validated a protocol for the rapid, culture-free, agnostic identification of pathogenic species from clinical canine samples, by cost-effective metagenomic sequencing. We have shown that this protocol is capable of detecting a wide array of species, representing over 90% of the urine and skin infections seen in the R(D)SVS HfSA. Although we did not test the remaining 10% of species due to their large number and relative rarity, our metagenomic extraction should effectively extract DNA from any species present in a sample, and we have shown up to 100% sensitivity and specificity in the identification of species from sequencing data alone. We intentionally developed a protocol that can also be adapted to a variety of other sample types, and the MagAttract protocol can be easily adapted for tissue, blood and other bodily fluids. In this way, one simple protocol can be deployed in a clinical setting to detect pathogens in a wide variety of infections, in different animals, in as little as 5 hours, compared to the 48 hours plus commonly seen in the current gold-standard diagnostics techniques.

Although the large amounts of host cell contamination we saw in some samples was problematic with regards to the prediction of AMR, a number of depletion techniques exist which could be incorporated into our protocol, at the cost of time and likely money, but with the benefit of allowing accurate AMR prediction from a greater range of samples. Outwith the samples with the highest levels of host DNA contamination, our ability to predict AMR was relatively accurate, approaching 95% in all but the most complicated co-infections. However, according to phenotypic AST a number of samples displayed intermediate levels of resistance to certain antimicrobials, mainly chloramphenicol, which were not predicted from the sequencing data. This suggests that as-yet unknown mechanisms may be responsible for intermediate resistance, or that certain mechanisms are currently missing from the AMR databases used (NCBI, ResFinder and CARD). Our current AMR prediction pipeline includes combining the results of three different tools (EPI2ME, Abricate and AMRFinderPlus), which function in different ways, in order to capture all potential information from our sequencing data. Future developments may include combining these tools into one easy-to-run workflow, as well as testing alternative tools and databases, such as the newly released, ISO-certified, abritAMR (55).

Lastly, we note that the flow cells (R9.4.1) and kits (SQK-RBK004) used here throughout development are now being replaced with R10.4.1 and SQK-RBK114. We expect these new flow cells and kits to incorporate seamlessly into our existing protocol and, indeed, they will likely improve the accuracy of the sequencing data produced, which could in turn improve the accuracy of our prediction of SNP-based AMR.

## Author statements

### Authors and contributors

Conceptualization: NR, AL, BW, GP, TN, RM, DG, JRF; Data curation: NR, AL; Formal analysis: NR; Funding acquisition: BW, GP, TN, DG, JRF; Investigation: AL, BW, NR; Methodology: NR, AL, BW; Project administration: JRF; Resources: GP, TN; Supervision: RM, DG, JRF; Validation: NR, AL, BW; Visualization: NR, BW; Writing - original draft: NR, JRF; Writing - review & editing: NR, AL, BW, GP, TN, RM, DG, JRF.

### Conflicts of interest

The authors declare there are no conflicts of interest affecting this work.

### Funding information

This work was funded by the Dog’s Trust Canine Welfare Grant, Direct Diagnostic Genomics Towards Effective Antimicrobial Use (“Dogstails”).

### Ethical approval

This work involved the use of non-experimental pet animals only and followed established internationally recognised high standards (‘best practice’) of individual veterinary clinical patient care. Samples were non-invasive and/or were excess from those taken during ordinary clinical treatment and, as such, were covered by the R(D)SVS’s ethical approval for research.

## Acknowledgements

Many thanks to Jennifer Harris, Easter Bush Pathology, R(D)SVS, University of Edinburgh, for sharing clinical samples and the results thereof.

The authors are grateful for the fantastic bioinformatics resource, CLIMB-BIG-DATA (developed by the MRC, grant number MR/T030062/1), without which much of the data analysis undertaken here would not have been possible.

For the purpose of open access, the authors have applied a Creative Commons Attribution (CC BY) licence to any Author Accepted Manuscript version arising from this submission.

## Notes

### Competing Interest Statement

The authors have declared no competing interest.

