## Supplementary Figures for "Rapid metagenomic sequencing for diagnosis and antimicrobial sensitivity prediction of canine bacterial infections"

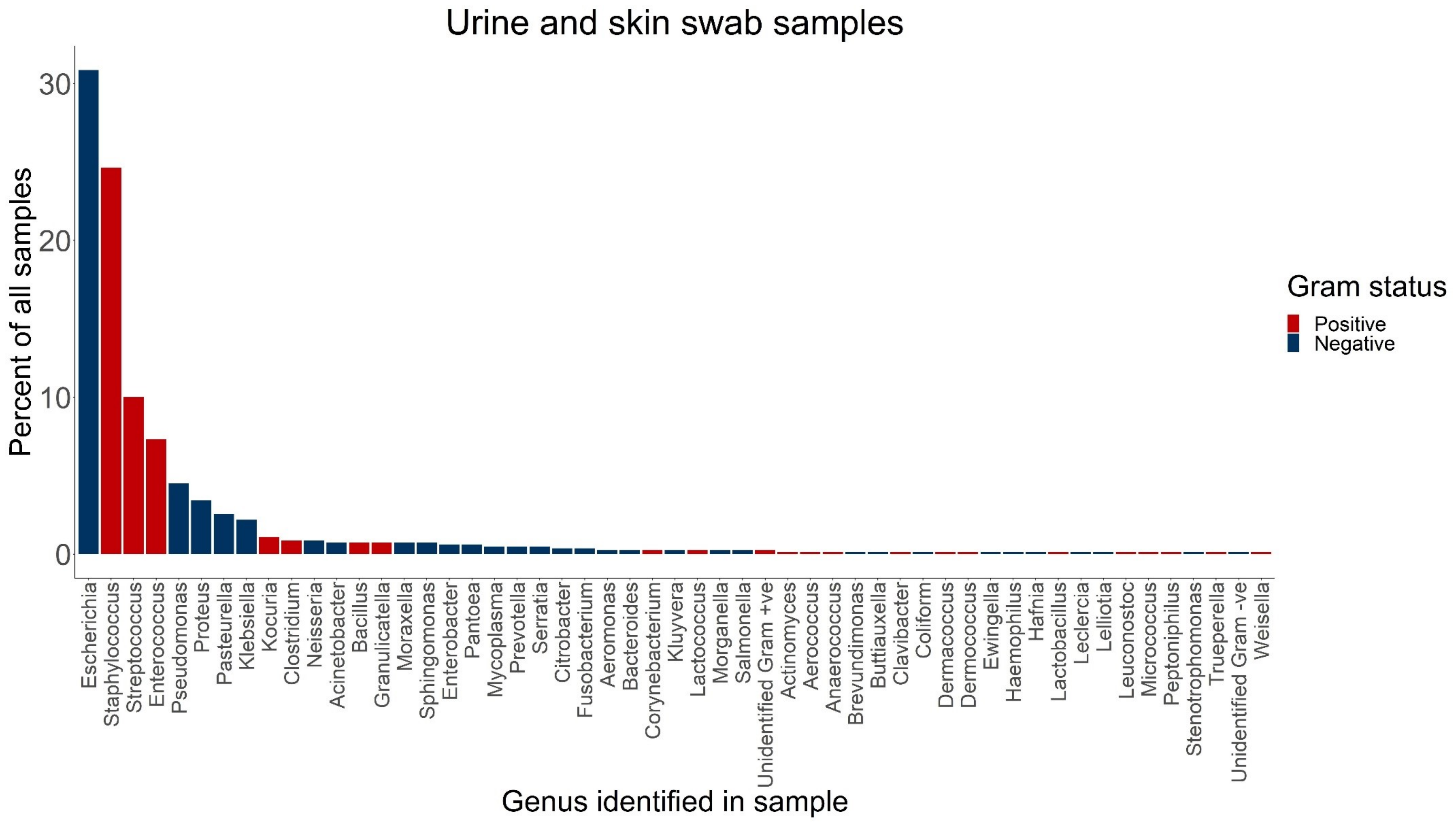


**Fig. S1 Pathogens identified in HfSA urine and skin swab samples, 2018 & 2019.**
Blue text indicates Gram –ve species, red text indicates Gram +ve species. There was roughly a 50:50 split (52% Gram –ve, 48% Gram-ve)

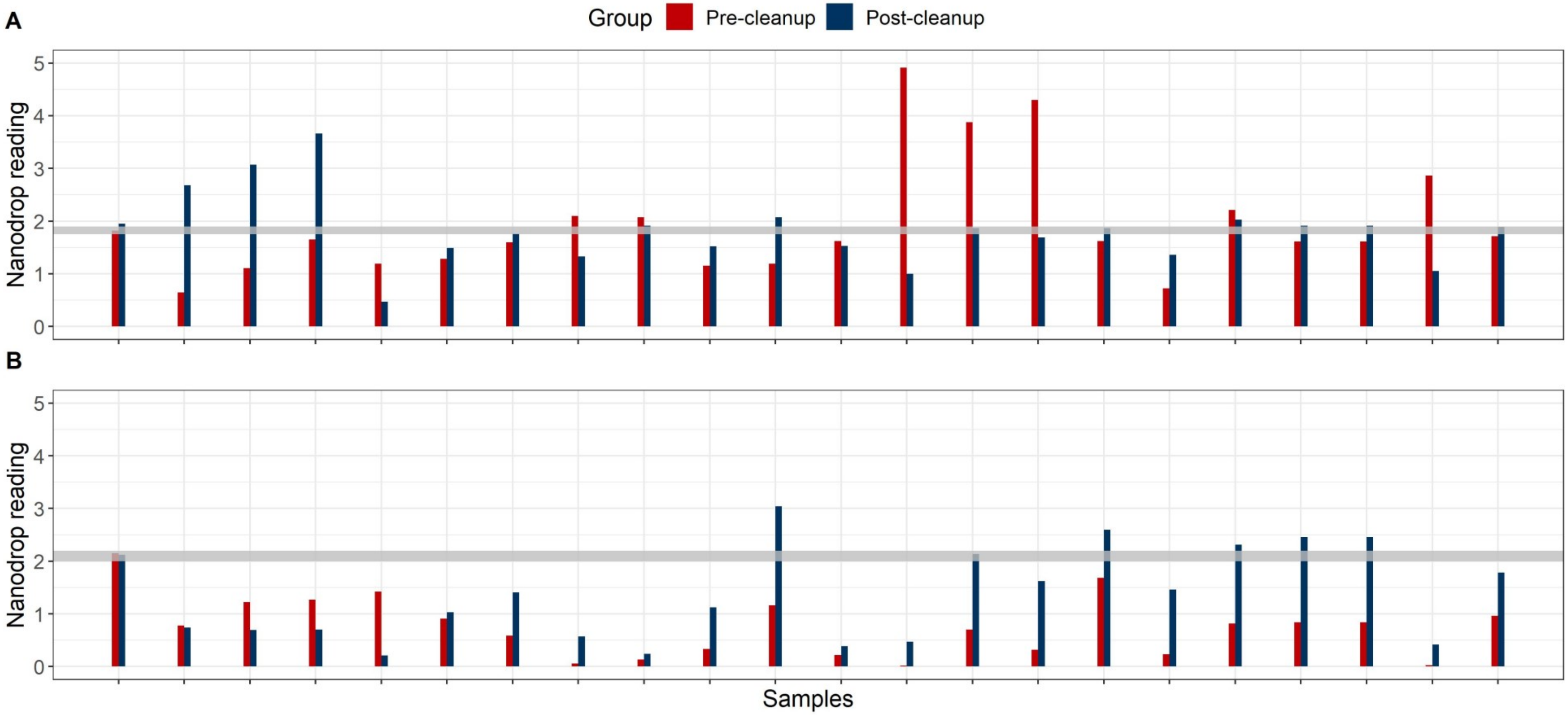


**Fig. 2 Nanodrop 260/280 (A) and 260/230 (B) ratios for 22 clinical samples, measured before and after ProNex bead clean-up.**
The dark grey rectangles on each plot indicate the ideal ranges for each ratio.

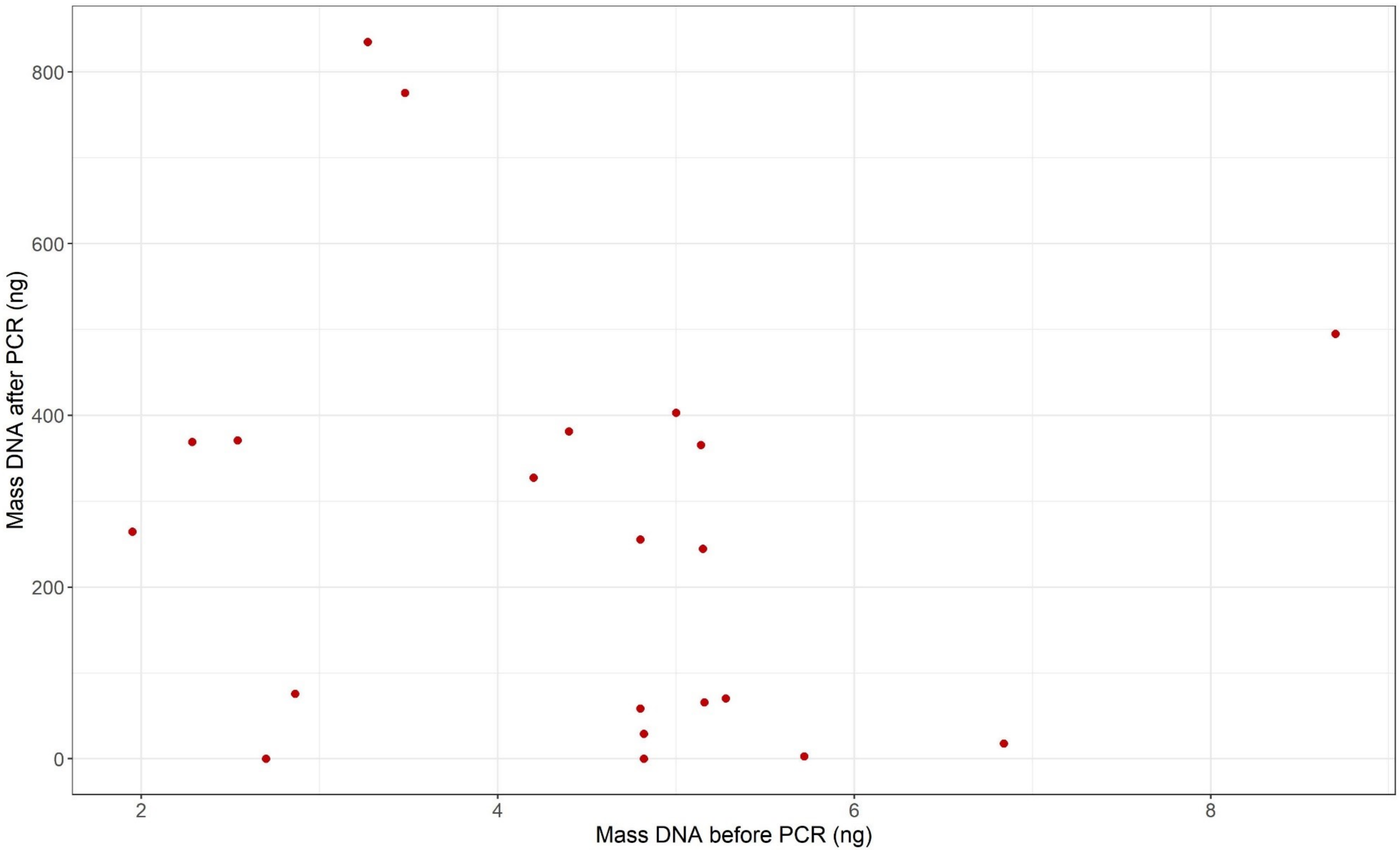


**Fig. S3 Inconsistent efficiency of the PCR in the Rapid PCR Barcoding (SQK-RPB004) library preparation kit.**
21 samples of varied starting concentration (hence varied mass of DNA used in PCR reaction) were amplified by the SQK-RPB004 PCR reaction, and their amplified DNA concentration in 10 µl was measured by the Qubit HS DNA kit.

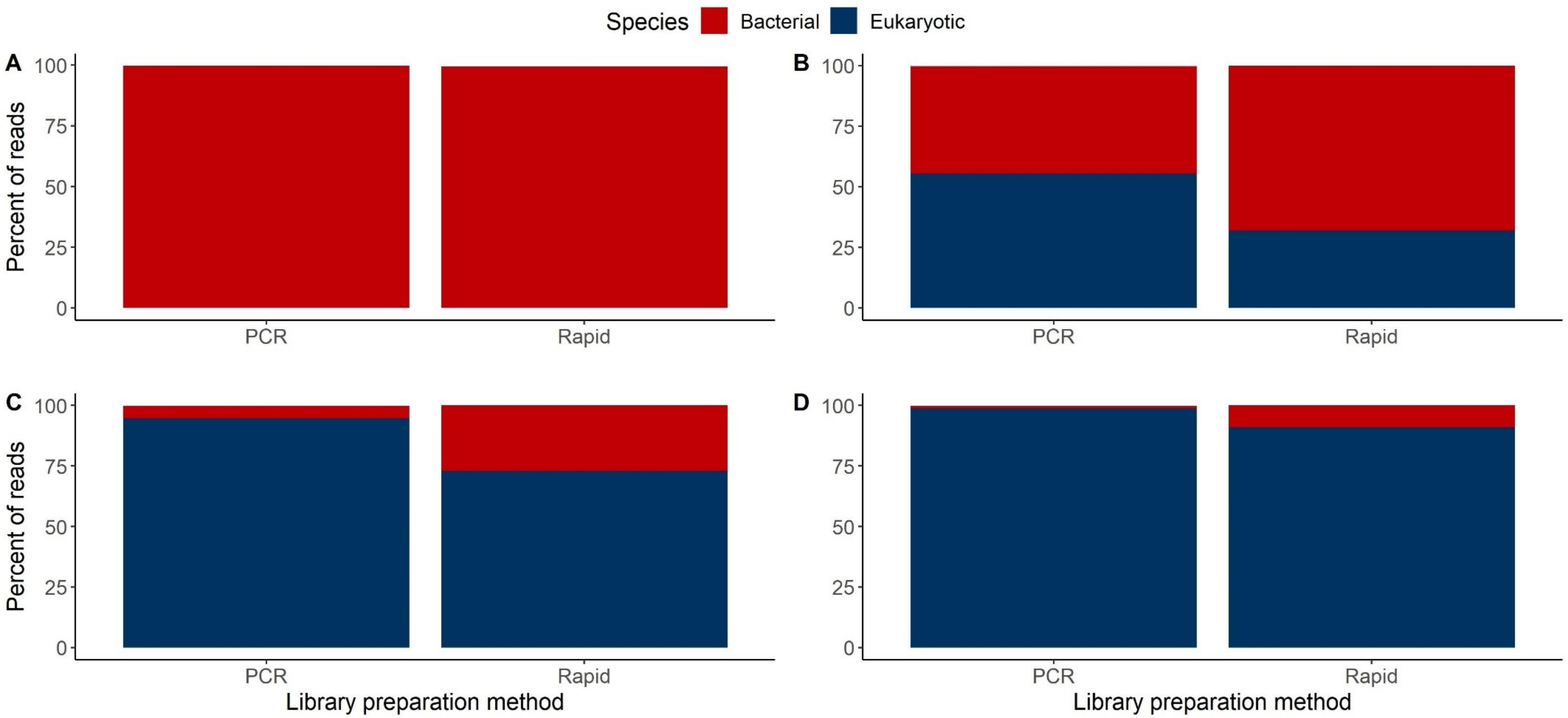


**Fig. S4 Selective eukaryotic DNA amplification by PCR.**
Four clinical samples were sequenced with both the Rapid PCR Barcoding (SQK-RPB004) and Rapid Barcoding (SQK-RBK004) kit for 24 hours on R9.4.1 MinION flow cells. A) and B) represent urine samples DTU09 and DTU16 respectively, while C) and D) skin swab samples SkSw08A and SkSw14 respectively.
